# Stromal HIF2 Regulates Immune Suppression in the Pancreatic Cancer Microenvironment

**DOI:** 10.1101/2021.05.21.445190

**Authors:** Yanqing Huang, Carolina J. Garcia Garcia, Daniel Lin, Nicholas D. Nguyen, Tara N. Fujimoto, Jun Zhao, Jaewon J. Lee, Vincent Bernard, Meifang Yu, Abagail M. Delahoussaye, Jae L. Phan, Amit Deorukhkar, Jessica M. Molkentine, Natividad R. Fuentes, Madeleine C. Turner, Dieter Saur, Anirban Maitra, Cullen M. Taniguchi

## Abstract

**Background & Aims:** Pancreatic ductal adenocarcinoma (PDAC) has a hypoxic, immunosuppressive stroma, which contributes to its resistance to immune checkpoint blockade therapies. The hypoxia-inducible factors (HIFs) mediate the cellular response to hypoxia, but their role within the PDAC tumor microenvironment remains unknown.

**Methods:** We used a dual recombinase mouse model to delete *Hif1α* or *Hif2α* in *α-smooth muscle actin* (*αSMA)*-expressing cancer-associated fibroblasts (CAFs) arising within spontaneous pancreatic tumors. The effects of CAF-*Hif2α* expression on tumor progression and composition of the tumor microenvironment were evaluated by Kaplan-Meier analysis, quantitative real-time polymerase chain reaction, histology, immunostaining, and by both bulk and single-cell RNA sequencing. CAF-macrophage crosstalk was modeled *ex vivo* using conditioned media from CAFs after treatment with hypoxia and PT2399, a HIF2 inhibitor currently in clinical trials. Syngeneic flank and orthotopic PDAC models were used to assess whether HIF2 inhibition improves response to immune checkpoint blockade.

**Results:** CAF-specific deletion of HIF2, but not HIF1, suppressed PDAC tumor progression and growth, and improved survival of mice by 50% (n = 21-23 mice/group, Log-rank *P* = 0.0009). Deletion of CAF-HIF2 modestly reduced tumor fibrosis and significantly decreased the intratumoral recruitment of immunosuppressive M2 macrophages and regulatory T cells. Treatment with the clinical HIF2 inhibitor PT2399 significantly reduced *in vitro* macrophage chemotaxis and M2 polarization, and improved tumor responses to immunotherapy in both syngeneic PDAC mouse models.

**Conclusions:** Together, these data suggest that stromal HIF2 is an essential component of PDAC pathobiology and is a druggable therapeutic target that could relieve tumor microenvironment immunosuppression and enhance immune responses in this disease.

## Introduction

Pancreatic ductal adenocarcinoma (PDAC) responds poorly to most cancer treatments, including immunotherapy^1^. This therapeutic recalcitrance may stem from PDAC’s extensive desmoplastic stroma, which suppresses anti-tumor immunity^2^ and increases intratumoral pressure^3^, resulting in severe hypoxia^4^ and impaired drug delivery^5^. Cancer-associated fibroblasts (CAFs) are the main components and producers of stroma in PDAC^6^. Efforts to physically disrupt the hypoxic stromal component through Sonic hedgehog protein inhibition^7^, selective fibroblast depletion^8^, or recombinant human hyaluronidase^9^ have effectively lowered stromal content but paradoxically led to worse outcomes in both preclinical studies and clinical trials. These data argue that the initially promising strategy of physically ablating the PDAC stroma may be clinically counterproductive, warranting a different approach.

The hypoxia-inducible factors 1 (HIF1) and 2 (HIF2) are stabilized in low oxygen and have been hypothesized to mediate therapeutic resistance^10^ and aggressive growth of PDAC^11^. Deletion of HIF1^12^ or HIF2^13^ in the pancreatic epithelial compartment failed to change overall survival in mice with spontaneous PDAC. However, the function of HIFs in other prominent compartments of the pancreatic tumor microenvironment (TME) remains unclear. Given the importance of the tumor stroma in PDAC oncobiology, we investigated the role of HIF signaling in CAFs and its impact on the PDAC TME.

Here, we elucidated the function of the HIFs within the PDAC stroma using a dual recombinase model to spatiotemporally alter HIF1 or HIF2 signaling only in activated fibroblasts reprogrammed within spontaneous murine pancreatic tumors (also known as CAFs). We found that CAF-specific deletion of HIF2, but not HIF1, improved survival from pancreatic cancer by reducing the recruitment of immunosuppressive macrophages. We further showed that therapeutic HIF2 inhibition improved responses to immune checkpoint blockade, indicating this is a potential combinatorial therapeutic strategy for PDAC.

## Materials and Methods

### Mice

All experimental mouse work adhered to the standards articulated in the Animal Research: Reporting of *In Vivo* Experiments guidelines. Additionally, all mouse work was approved by the Institutional Animal Care and Use Committee of The University of Texas MD Anderson Cancer Center. Both female and male mice were used in this study. Mice were maintained on a 12-hour light/dark cycle and were provided with sterilized water and either standard rodent chow (Prolab Isopro RMH 3000 irradiated feed) or a tamoxifen diet (Teklad, TD.130855, 250 mg tamoxifen/kg). Experiments were carried out during the light cycle.

*FSF-Kras*^*G12D/+*^;*P53*^*frt/frt*^ mice were gifts from Dr. David Kirsch (Duke University)^14, 15^. *Pdx1*^*Flp/+*^ mice were gifts from Dr. Dieter Saur (Technical University, Munich)^16^. *αSMA*^*CreERT2/+*^ mice were gifts from Dr. Richard Premont (Case Western Reserve University)^17^. *Hif1α* ^*fl/fl*^ (RRID:IMSR_JAX:007561), *Hif2α* ^*fl/fl*^ (RRID:IMSR_JAX:008407), *LSL-tdTomato* (RRID:IMSR_JAX:007914), and C57BL/6 (RRID:IMSR_JAX:000664) mice were obtained from Jackson Laboratories. *FSF-Kras*^*G12D/+*^;*P53*^*frt/frt*^ mice were bred with *Pdx1*^*Flp*^ mice to produce *FSF-<u>K</u>ras*^*G12D/+*^;*<u>P</u>53*^*frt/frt*^;*Pdx1*^<u>*F*</u>*lp*^ (KPF) mice. KPF mice were bred with *αSMA*^*CreERT2/+*^ mice and their progeny was bred with *Hif1α* ^*fl/fl*^ and *Hif2α* ^*fl/fl*^ mice to produce KPF CAF-HIF1 and KPF CAF-HIF2 mice, respectively. *LSL-<u>K</u>ras*^*G12D/+*^;*Tr<u>p</u>53*^*fl/fl*^;*Ptf1a*^<u>*C*</u>*re*/+^ (KPC) mice and *EGLN1/2/3*^*fl/fl*^ mice were previously bred and backcrossed to C57BL/6 mice in our lab^18, 19^. Genotyping was performed as described previously^20^. Littermate controls were used in all experiments.

### Isolation and *Ex Vivo* Analysis of Fibroblasts and CAFs

tdTomato reporter mice were bred with *αSMA*^*CreERT2/+*^;*Hif2α* ^*fl/fl*^ mice to produce *αSMA*^*CreERT2/+*^;*Hif2α* ^*fl/fl*^;*tdTomato*^*LSL/LSL*^ mice. Normal pancreata from the tdTomato progeny and from *EGLN1/2/3*^*fl/fl*^ mice and whole tumors from KPC mice were minced and digested with 1 mg/mL Collagenase V (Sigma, C9263-500MG) for 30 minutes at 37°C and 130 rpm/min followed by digestion with TrypLE (Thermo Fisher Scientific, 12605036) for 10 minutes at 37°C. Cells were seeded in T175 flasks with DMEM (ATCC, 30-2002) plus 10% (v/v) FBS (MilliporeSigma, F4135) and 1% Pen/Strep. Upon reaching 70% confluence, cells were passaged and incubated at 37°C for 30 minutes before the media was refreshed. The attached cells became enriched for fibroblasts or CAFs after 2-5 passages. Normal fibroblasts were immortalized with a pBABE-hydro-hTERT lentivirus (Addgene, #1773).

*Hif2α* ^*fl/fl*^; *αSMA*^*CreERT2/+*^; *tdTomato*^*LSL/LSL*^ fibroblasts were treated with DMSO, 4-hydroxytamoxifen (4-OHT), or Adeno-Cre as a positive control. Fibroblasts were then genotyped as described previously^20^ using the primers listed in Supplementary Table 3 and imaged with an Olympus FV500 laser scanning confocal microscope (Olympus USA).

### Histopathology and Immunohistochemistry

Spontaneous PDAC tumors were harvested from KPF-CAF HIF2 wild-type (WT) and knockout (KO) mice and fixed with 10% neutral buffered formalin, subjected to ethanol dehydration, washed in Histoclear (National Diagnostics, HS2001GLL), and embedded in paraffin. Then 5-µm-thick tissue slices were cut, mounted onto slides, and stained with H&E. Masson’s trichrome staining was performed in the Research Histology Core Lab (RHCL) at MD Anderson. Histopathologic assessment of H&E staining and fibrosis scoring of trichrome-stained KPF-CAF HIF2 tumor slides were performed by a pathologist (J. Zhao) who was blinded to genotype. Immunohistochemistry (IHC) was performed as previously described ^18^ using anti-HIF2α (1:200, Abcam Cat# ab199, RRID:AB_302739), anti-F4/80 (1:200, Abcam Cat# ab6640, RRID:AB_1140040), and anti-FoxP3 (1:200, Abcam Cat# ab20034, RRID:AB_445284) antibodies.

### Epithelial HIF1 and HIF2 KO

We isolated epithelial PDAC cells from KPF-HIF1^fl/fl^ or KPF-HIF2^fl/fl^ mice and induced recombination *ex vivo* via infection with Adeno-Cre or control Adeno-GFP. Recombined KPF cells were resuspended in PBS and Matrigel in a 1:1 ratio and orthotopically implanted into the pancreata of immunocompromised mice.

### Bulk RNA Sequencing

Frozen tumors from KPF-CAF HIF2 WT and KO mice were homogenized and RNA was purified using an RNeasy mini kit (QIAGEN, 74106). Library preparation and sequencing were performed in the Sequencing and Microarray Facility at MD Anderson. RSEM software package (RRID:SCR_013027) was used to quantitate transcript abundance from RNA-seq data^21^. Differential expression analysis was performed using DESeq2 software package (RRID:SCR_015687). Supplementary Table 1 shows the normalized transcript counts. Gene Set Enrichment Analysis was performed to identify significantly enriched pathways (FDR < 0.15).

### CAF Conditioned Media Harvest

CAFs isolated from KPC tumors and immortalized normal fibroblasts isolated from EGLN1/2/3^fl/fl^ mice were seeded in DMEM with 10% FBS and 1% Pen/Strep at 5 x10^5^ density in 60-mm cell plates and cultured overnight. The media was replaced with DMEM containing 0.5% FBS, and cells were transferred to a hypoxia chamber (InvivO2, Baker Ruskinn) set at 1% O_2_ and treated with increasing concentrations of PT2399 (Peloton Therapeutics/Merck) for 48 hours ^22^. Cell media was collected and centrifuged at 3,000 rpm for 5 minutes, and the supernatant was stored as conditioned media at −80°C until the experiment.

### Macrophage Transwell Migration Assay

Authenticated RAW 264.7 murine macrophages were purchased from ATCC (Cat# TIB-71, RRID:CVCL_0493) and maintained in DMEM with 10% FBS at 37°C with 5% CO_2_. Cell suspension aliquots were reseeded into new culture vessels and early passages were used for all experiments. Macrophage migration was tested in 24-well Transwell permeable plates with 8-μm-pore polycarbonate membrane inserts (Corning, 3422). Macrophages were resuspended in 100 μL of culture media and added to the upper chamber, while 600 μL of conditioned media from CAFs or normal fibroblasts was added to the lower chamber as a chemoattractant. Cells were allowed to migrate through the membrane insert for 24 hours and nonmigrating macrophages were removed with a cotton swab. Migrated macrophages were fixed and stained with hematoxylin for imaging using a light microscope (Leica DMi1). At least 3 random nonoverlapping fields (10x magnification) were quantified using ImageJ software (RRID:SCR_003070).

### Quantitative Real Time-Polymerase Chain Reaction

Frozen tumors from KPF-CAF HIF2 WT and KO mice were homogenized and RNA was purified using an RNeasy mini kit (QIAGEN, 74106), and reverse transcription was performed with the QuantiTect reverse transcription kit (QIAGEN, 205313). Quantitative real-time polymerase chain reaction (qRT-PCR) was carried out using a QuantiFast SYBR Green PCR kit (QIAGEN, 204056) on a StepOnePlus real-time PCR system (Applied Biosystems). qRT-PCR analysis of *Arg1* was also performed on RAW 264.7 murine macrophages treated with the indicated conditions. Primers are listed in Supplementary Table 3.

### Single-Cell RNA Sequencing

Single-cell suspensions were prepared by mincing KPF CAF-HIF2 WT and KO tumors, digesting them with 0.5 mg/mL Liberase (Sigma, LIBTH-RO 5401135001) for 30 minutes at 130 rpm, and passing them through a 100-μm cell strainer. Samples were then incubated with Accutase (Sigma, A6964) for 10 minutes at 37°C in a shaker, followed by treatment with ACK lysing buffer (ThermoFisher, A1049201) to eliminate erythrocytes. Samples were filtered through a 30-μm cell strainer and single cells were resuspended in PBS (GE Healthcare Life Sciences, SH30256.01) with 0.1% BSA. Cell viability was measured using Trypan Blue (Bio-Rad, 1450021). Single-cell suspensions were loaded into a 10x Genomics Chromium instrument to generate gel beads in emulsion. Approximately 5,000 cells were loaded per channel. Single-cell complementary DNA (cDNA) libraries were prepared using a Chromium Single Cell 3’ Library & Gel Bead kit v2 (10x Genomics, PN-120237) and sequenced using a NextSeq 500 (Illumina). The mean number of reads per cell was approximately 25,000 and the median number of genes detected per cell was approximately 2,000.

The raw data were processed using cellranger count (Cell Ranger v2.1.1, 10x Genomics) based on the mm10 mouse reference genome. Subsequent data analysis was done in R using the Seurat package v3.0 (RRID:SCR_007322) and default parameters. Uniform Manifold Approximation and Projection (UMAP) dimensionality reduction and graph-based clustering of cells were performed, and clusters were assigned to cell populations using known signature genes. Supplementary Table 2 shows the genes enriched in each cell population versus all other cells.

### Immunotherapy Experiments

We obtained KPC cells from Dr. Anirban Maitra that were authenticated by short tandem repeat profiling and were confirmed to be *Mycoplasma* free by real-time PCR (CellCheck Mouse 19 Plus, IDEXX Laboratories, Inc.). For the flank model, 1 × 10^6^ KPC cells were resuspended in PBS and Matrigel (Corning) in a 1:1 ratio and subcutaneously implanted into the right flanks of syngeneic 10-week-old C57BL/6 female mice. Murine αCTLA4 (BioXCell, BE0164) or isotype control was administered intraperitoneally (IP) every 3-4 days at 250 μg/mouse, beginning 13 days after implantation (Figure 4G). PT2399 was resuspended in 10% ethanol, 30% PEG400, and 60% methylcellulose/water/Tween 80 and administered 5 days per week (Monday-Friday), twice daily, at 50 mg/kg via oral gavage. Treatments lasted 2 weeks, and tumor dimensions were measured with a caliper to calculate approximate volumes.

For the orthotopic model, 2 × 10^5^ KPC cells were resuspended in PBS and Matrigel in a 1:1 ratio and injected into the tail of the pancreas of syngeneic 12-week-old C57BL/6 male mice. After 2 weeks of recovery, murine αCTLA4 (clone 9D9, Merck) and murine αPD1 (muDX400, Merck) or isotype control were administered IP every 4 days at 20 μg/mouse, 200 μg/mouse, and 220 μg/mouse, respectively for 2 weeks. PT2399 was administered 5 days per week, twice daily for 3 weeks, at 50 mg/kg via oral gavage (Figure 4I). Tumor burden was monitored by ultrasound. Mice were age-matched but group assignment was unblinded.

### Statistical Methods

Survival was analyzed by the Kaplan-Meier method and log-rank test. Student’s *t* test was used to analyze parametric data sets and the Mann–Whitney *U* test was used for non-parametric data sets. All statistical analyses were performed using GraphPad Prism V.8 (RRID:SCR_002798), with a significance level of α = 0.05.

## Results

### Deletion of Stromal HIF2 Delays PDAC Progression and Enhances Survival

We used a dual recombinase system to constrain the deletion of *Hif1α* or *Hif2α* to CAFs within autochthonous PDAC tumors. Mice with FlpO-responsive alleles of both oncogenic *Kras* (*FSF*-***K****ras*^*G12D/+*^)^14^ and homozygous *Trp53* (*Tr****p****53*^*frt/frt*^)^15^ were crossed with mice expressing FlpO in pancreatic tissue (*Pdx1-* ***F****lpO*)^16^ to generate KPF mice. These mice developed spontaneous PDAC over a timeframe and with a penetrance similar to those in KPC mice (Cre-driven model), and both models recapitulate human PDAC^16^. KPF mice were subsequently bred with mice harboring conditional null alleles of *Hif1α* (*Hif1α*^*fl/fl*^)^23^ or *Hif2α* (*Hif2α*^*fl/fl*^)^24^, driven by expression of the *Cre-ER*^*T2*^ transgene under the control of the *α-smooth muscle actin* (*αSMA*, also known as *Acta2*) promoter which marks CAFs (Figure 1A)^17^. We confirmed deletion of HIF1 or HIF2 through *ex vivo* analyses of activated fibroblasts isolated from tdTomato reporter mice (Supplementary Figure 1A-B). Once weaned, mice were fed normal chow or tamoxifen chow to generate KPF CAF-HIF WT and KPF CAF-HIF KO mice, respectively (Figure 1B). Mice were screened for tumors weekly by ultrasound. The median age at tumor onset was 10.3 weeks (range: 7.1–21.1) in both the

**Figure 1.**
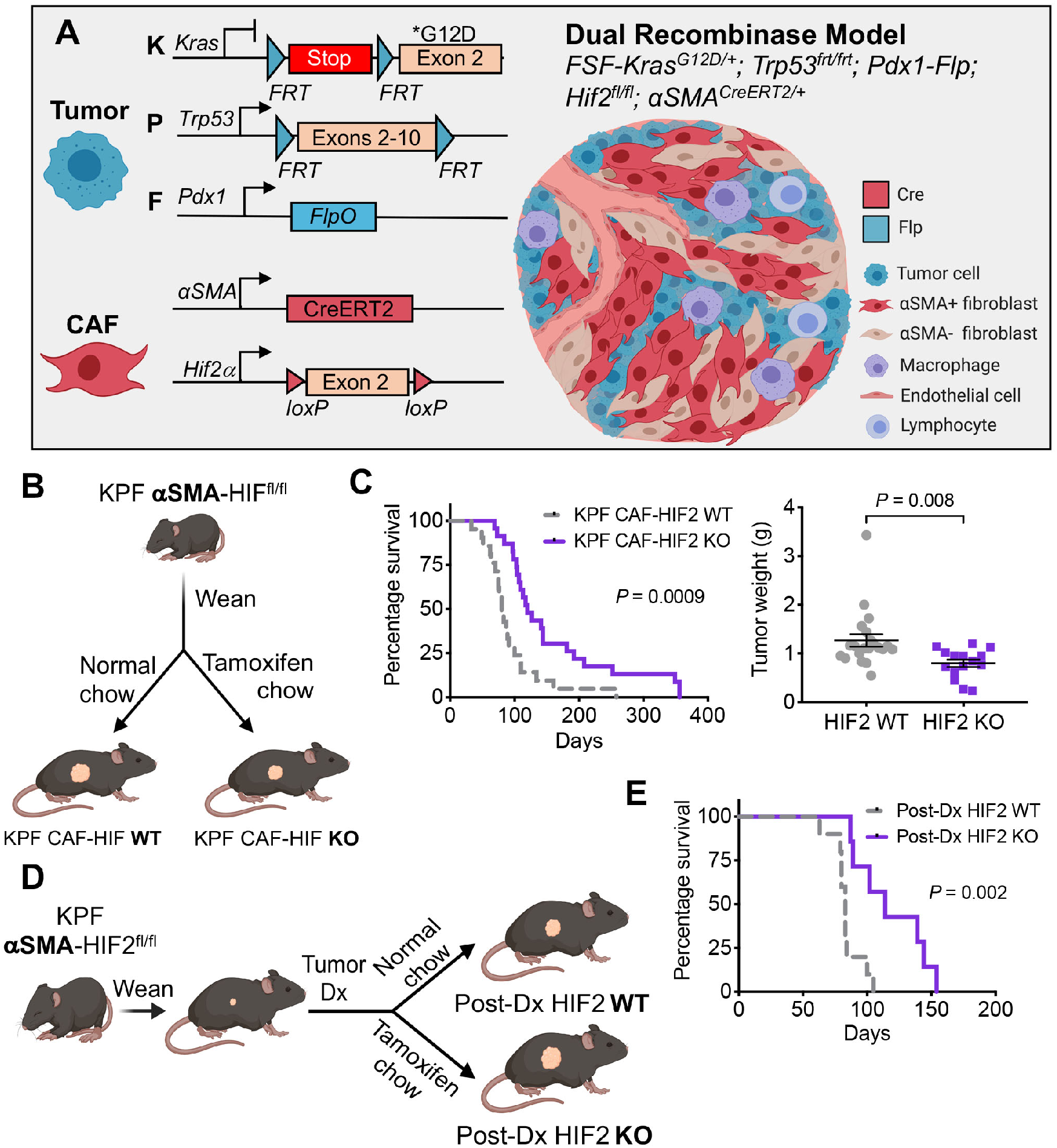
Deletion of stromal HIF2 delays PDAC progression and enhances survival. *(A)* Dual recombinase genetic strategy to develop a PDAC model with HIF1 or HIF2 knockout (KO) in *αSMA*^*+*^ cells in a tamoxifen-induced manner using KPF mice. *(B)* Experimental design to generate KPF CAF-HIF wildtype (WT) control and KPF early CAF-HIF KO mice. *(C) Left*: Kaplan-Meier curves showing percentage survival for KPF CAF-HIF2 WT (n = 21) and KO (n = 23) mice. *P*, by log-rank test. *Right*: Tumor weights of KPF CAF-HIF2 WT (n = 21) and KO (n = 15) mice. Mean ± SEM; *P*, by Student’s *t* test. *(D)* Experimental design to generate post-diagnosis (post-Dx) KPF CAF-HIF2 WT and KO mice. *(E)* Kaplan-Meier curves showing percentage survival for post-Dx KPF CAF-HIF2 WT (n = 10) and KO (n = 7) mice. *P*, by log-rank test. See also Supplementary Figures 1-3.

WT and KO groups. Immunohistochemical analysis confirmed a reduction of HIF2 expression in CAFs within KPF tumors (Supplementary Figure 1C). Surprisingly, loss of stromal HIF1 had no effect on tumor growth or survival (median survival, 91 days for KO versus 100 days for WT; Supplementary Figure 1D-E).

In contrast, HIF2 ablation in CAFs significantly decreased tumor growth and improved survival (median survival, 120 days for KO versus 80 days for WT; n = 21-23 mice/group, Log-rank *P* = 0.0009; Figure 1C). Histological analyses of the pancreata revealed well-differentiated PDAC foci in both groups, yet remarkably, we found no gross or microscopic evidence of tumor tissue in the sections analyzed from six of the HIF2-depleted mice, suggesting that deletion of stromal HIF2 may also influence PDAC oncogenesis and/or progression (Supplementary Figure 2A-C)^13^. Importantly, there were no statistically significant differences in tumor fibrosis associated with the presence or absence of stromal HIF2 (n = 8-12 tumors/group, *P* = 0.0506; Supplementary Figure 2D)^25^.

We next assessed how stromal HIF2 deletion after PDAC onset impacted survival, as this would more closely reflect the timeline of therapeutic HIF2 targeting in patients. We generated another cohort of KPF αSMA-HIF2^fl/fl^ mice and fed them normal chow or tamoxifen chow after tumors were diagnosed by ultrasound (Figure 1D). This late abrogation of HIF2 in CAFs still improved survival by 37.3% compared to the survival of control mice (median, 114 days versus 83 days; *P* = 0.002; Figure 1E), which was similar to the median survival of mice receiving tamoxifen chow at weaning. Together, these data suggest that stromal HIF2, but not HIF1, plays a critical role in PDAC development and progression.

To confirm that this survival advantage was mediated by HIF2 depletion in CAFs, and not in tumor cells, we isolated cancer cells from KPF tumors with *Hif1α*^*fl/fl*^ or *Hif2α*^*fl/fl*^ alleles and induced *ex vivo* recombination by infection with Cre or control GFP adenovirus. These KPF cells were orthotopically implanted into the pancreata of immunocompromised mice (Supplementary Figure 3A). We found that deletion of HIF1 or HIF2 in tumor cells had no impact on tumor growth (Supplementary Figure 3B-C), confirming cell non-autonomous functions of HIF in PDAC.

### Stromal HIF2 Regulates Macrophage Recruitment to PDAC Tumors

We performed bulk RNA sequencing (RNA-seq) to understand the mechanism by which loss of HIF2 in CAFs suppressed tumor growth. Transcriptomic analysis revealed a stromal HIF2-dependent immune gene signature with enrichment in multiple pathways related to myeloid/macrophage biology (Figure 2A, Supplementary Figure 4A-B, and Supplementary Table 1). Deletion of HIF2 in CAFs led to downregulation of genes involved in macrophage migration, differentiation, and activation, including *Mmp9, Cd74, Tgfb1*, and *Itgam*; these results were validated by qRT-PCR (Figure 2B-C and Supplementary Figure 4C). We next compared tumor-associated macrophage (TAM) infiltration by F4/80 IHC and observed significantly fewer TAMs in KPF CAF-HIF2 KO tumors than in controls (n = 5 tumors/group, *P* = 0.028; Figure 2D). These results suggest that HIF2 signaling in CAFs regulates macrophage recruitment to PDAC tumors.

**Figure 2.**
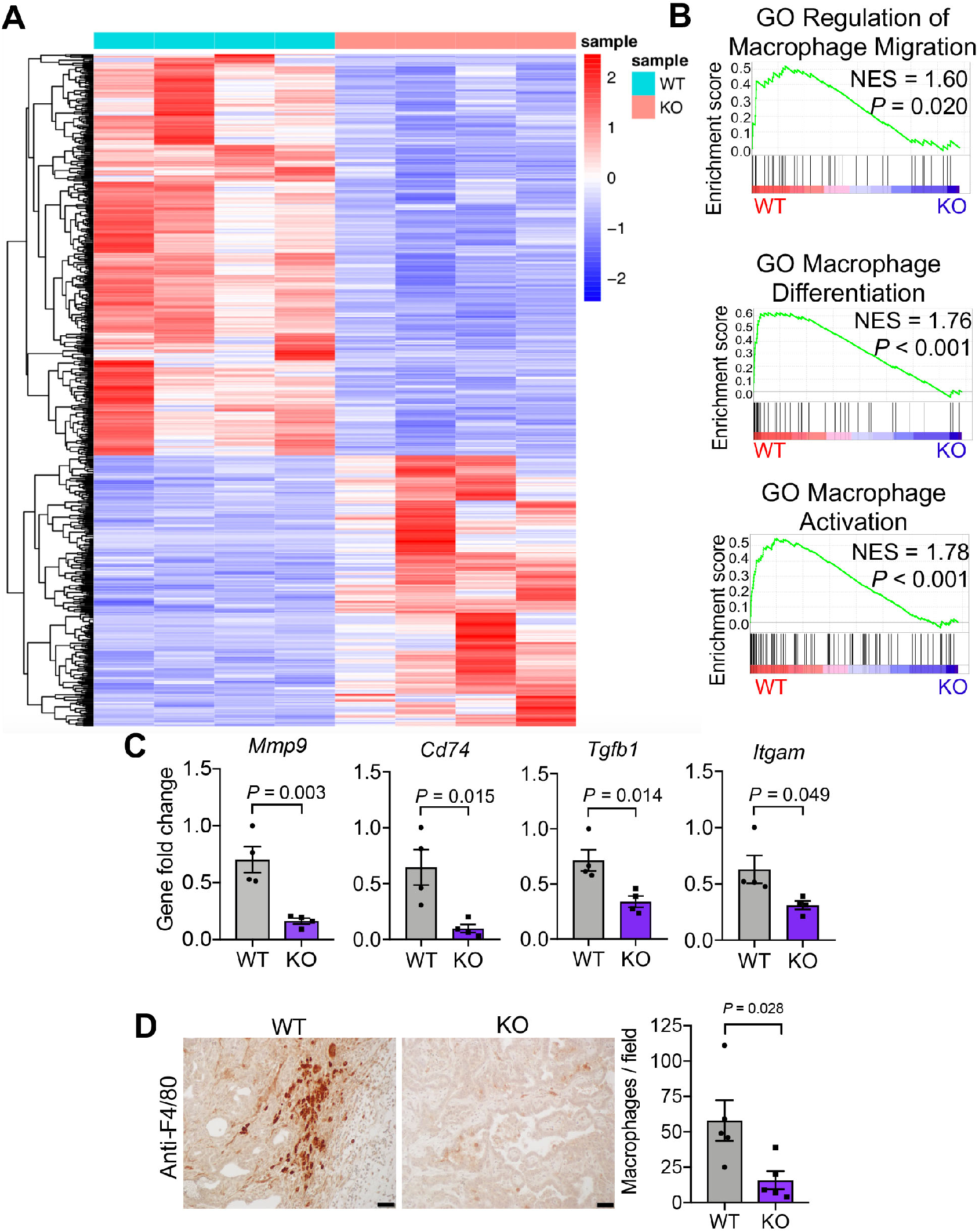
Stromal HIF2 regulates macrophage recruitment to PDAC tumors. *(A)* Heatmap of the top expressed genes using bulk RNA-seq data from KPF CAF-HIF2 tumors (n = 4/group). *(B)* Gene set enrichment analysis of tumors in (A) correlates CAF-HIF2 function with macrophage migration, differentiation, and activation. GO, gene ontology; NES, normalized enrichment score. *(C)* qRT-PCR confirmed the downregulation of genes involved in the pathways in *(B). (D) Left:* Representative IHC images of CAF-HIF2 tumors stained for F4/80 (n = 5/group). Scale bar, 50 µm. *Right:* Quantification of F4/80+ macrophages per field. All error bars represent mean ± SEM and each dot denotes a biological replicate. *P*, by Student’s *t* test. See also Supplementary Figure 4 and Supplementary Table 1.

To evaluate this hypothesis, we established CAF and normal fibroblast lines from spontaneous pancreatic tumors and normal pancreata, respectively. Both cell lines were cultured in hypoxia to stabilize HIF2 and to approximate *in vivo* TME conditions, and were then treated with either vehicle or the clinical HIF2 inhibitor PT2399^22^. We found that conditioned media from hypoxic CAFs stimulated macrophage migration in a HIF2-dependent fashion (Figures 3A-B). Stimulation of macrophage migration by CAFs appears to be specific to fibroblasts reprogrammed in the PDAC TME, as fibroblasts isolated from normal pancreata lacked the ability to stimulate macrophage migration (Supplementary Figure 5A). These results strongly suggest that a HIF2 coordinates CAF-TAM crosstalk in a paracrine fashion.

**Figure 3.**
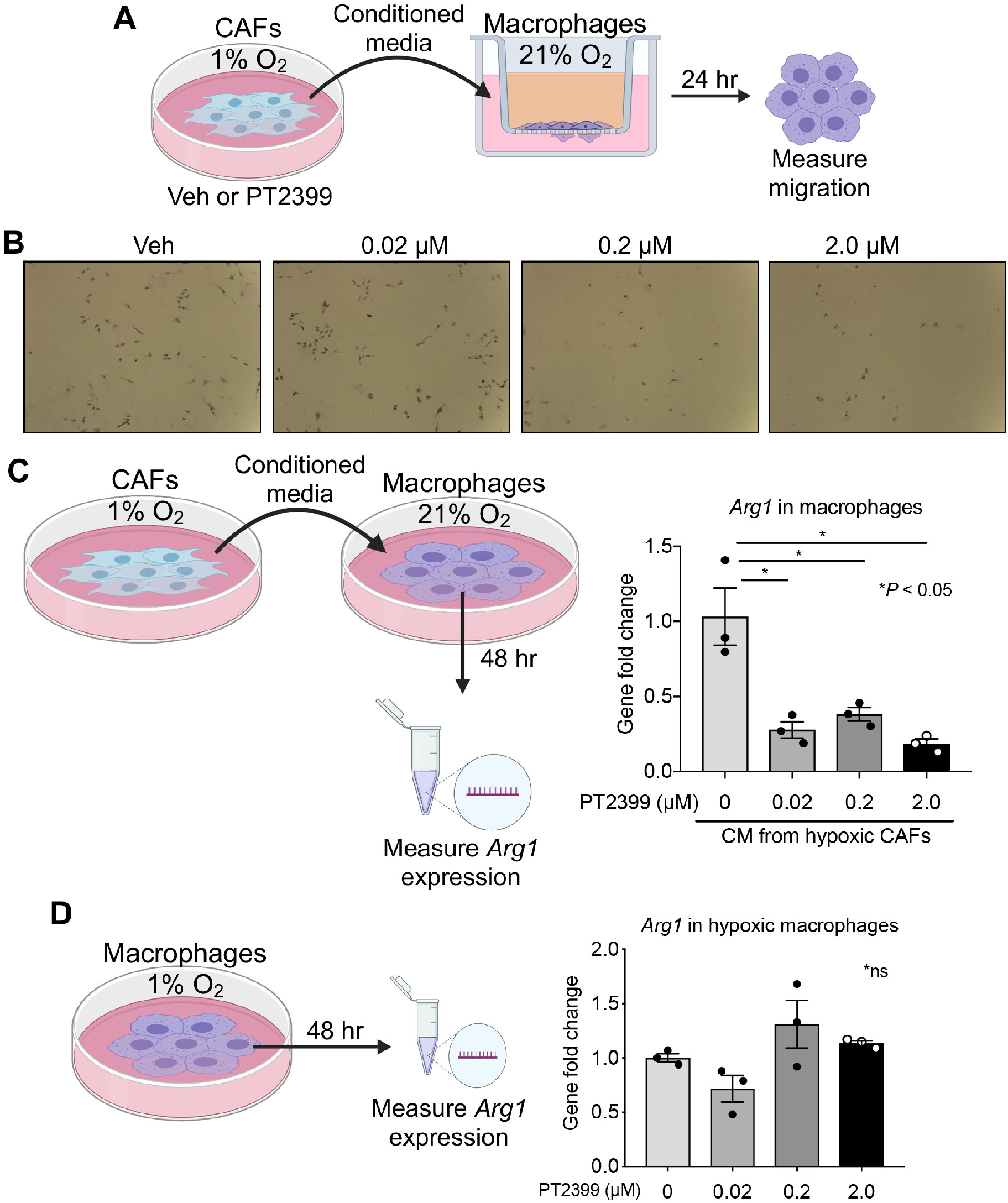
Hypoxic CAFs promote macrophage M2 polarization in a HIF2-dependent paracrine fashion. *(A)* Macrophages were cultured in transwell plates and incubated with conditioned media (CM) collected from hypoxic CAFs treated with vehicle (veh) or PT2399. *(B)* Representative bright-field images of the transwell assay (10x). *(C)* Macrophages were incubated with CM collected from hypoxic CAFs treated with veh or PT2399 and *Arg1* expression was measured by qRT-PCR. *(D)* Hypoxic macrophages were directly treated with veh or PT2399 and *Arg1* expression was measured by qRT-PCR. All error bars represent mean ± SEM; *P*, by Student’s *t* test. See also Supplementary Figure 5.

### Hypoxic CAFs Promote Macrophage M2 Polarization in a HIF2-Dependent Paracrine Fashion

Macrophages are functionally classified as either M1, which are classically activated and pro-inflammatory, or M2, which are alternatively activated during the resolution phase of inflammation, and thus display an immunosuppressive phenotype^26^. In many cancers, including PDAC, M2 macrophages are associated with worse outcomes because they promote metastasis and suppress anti-tumor immune responses via the expression of checkpoint ligands and by induction of regulatory T cells (Tregs)^26, 27^. CAFs have been linked to M2 repolarization of TAMs in PDAC^27^, yet the roles of hypoxia and HIF2 in this context remain unclear.

To understand whether HIF2 signaling in CAFs drives macrophage M2 repolarization, we stimulated murine macrophages with conditioned CAF media and assessed expression of *Arg1*, an M2 polarization marker (Figure 3C). We found that hypoxia, and therefore HIF2 expression, increased *Arg1* levels by 4-fold compared to controls (Supplementary Figure 5B). Moreover, HIF2 inhibition via PT2399 in hypoxic CAFs impaired the ability of conditioned media from these cells to induce M2 polarization, indicating that the paracrine CAF signal is HIF2-dependent (Figure 3C). Conditioned media from hypoxic normal pancreatic fibroblasts failed to induce M2 polarization (Supplementary Figure 5C), confirming that stimulation of macrophages by CAFs is specific to fibroblasts reprogrammed in the PDAC TME. Furthermore, direct HIF2 inhibition in macrophages using PT2399 did not affect M2 polarization (Figure 3D). Taken together, these findings support the notion that hypoxic CAFs activate TAMs in a HIF2-dependent paracrine fashion.

Vascular endothelial growth factor (VEGF) is a potent immunosuppressive factor known to induce M2 repolarization in TAMs^28^. Since *Vegf* is a hypoxia-inducible gene^29^, we measured *Vegf* expression in hypoxic CAFs treated with PT2399 (Supplementary Figure 5D), and found no differences, indicating that *Vegf* is not a critical component of the HIF2 regulation of immunosuppression by CAFs in our model.

### Deletion of Stromal HIF2 Reduces PDAC Immunosuppression

We performed single-cell RNA sequencing (scRNA-seq) to interrogate the impact of CAF-specific HIF2 signaling on other cells in the PDAC TME. We analyzed the transcriptomes from 22,635 single cells isolated from three KPF

CAF-HIF2 WT tumors (10,703 cells) and three KPF CAF-HIF2 KO tumors (11,932 cells). Graph-based clustering of cells after UMAP dimensionality reduction identified 32 clusters that were assigned to seven major cell types using signature genes (Figure 4A, Supplementary Figure 6A-C, and Supplementary Table 2). All of the cell populations identified were represented in both experimental groups and in all six mice, with 52.9% of the cells analyzed being identified as epithelial/tumor cells, 22.6% as myeloid cells, 19.8% as fibroblasts, and the remaining cells as endothelial cells, B cells, neutrophils, and T cells (Figure 4A and Supplementary Figure 6D).

**Figure 4.**
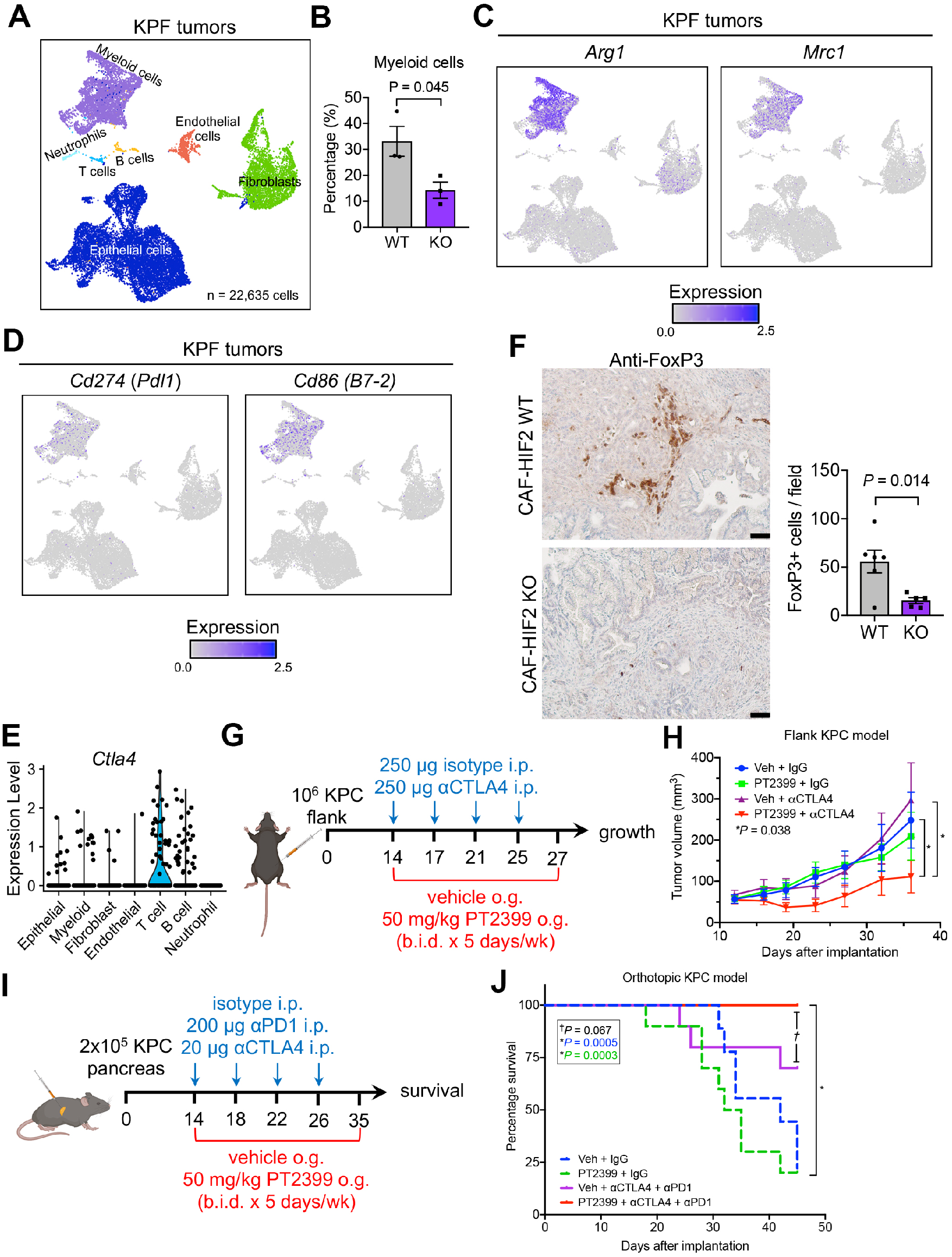
Inhibition of HIF2 signaling in CAFs reduces PDAC immunosuppression and enhances response to checkpoint immunotherapy. *(A)* UMAP of scRNA-seq analysis of 22,635 cells isolated from KPF CAF-HIF2 WT tumors (10,703 cells; n = 3 mice) and KPF CAF-HIF2 KO tumors (11,932 cells; n = 3 mice). Cell types were identified through graph-based clustering followed by manual annotation using marker genes. *(B)* Percentage of myeloid cells in each tumor. *(C)* M2-polarized TAMs were identified within the myeloid cell population via expression of *Arg1* and *Mrc1. (D)* Immunosuppressive TAMs were identified within the myeloid cell population via expression of *Cd274* (*Pdl1*) and *Cd86* (*B7-2*). *(E)* Violin plots showing findings on scRNA-seq analysis of *Ctla4* in KPF CAF-HIF2 WT and KO tumors in all identified cell types. *(F) Left:* Representative IHC images of CAF-HIF2 WT and KO tumors stained for FoxP3 (n = 5-6/group); scale bars, 50 µm. *Right:* Quantification of FoxP3+ Tregs per field. *(G)* Schematic for administration of PT2399 + αCTLA4 in a syngeneic flank KPC model. i.p., intraperitoneal; o.g., oral gavage; b.i.d., *bid in die* (twice a day). *(H)* Tumor growth curve from *(A)* (n = 10/group). Veh, vehicle; *P*, by Mann–Whitney *U* test. *(I)* Schematic for administration of PT2399 + αCTLA4/αPD1 in a syngeneic orthotopic KPC model. *(J)* Kaplan-Meier curves showing percentage survival for *(C)* (n = 10/group); *P*, by log-rank test. All error bars represent mean ± SEM; *P*, by Student’s *t* test unless otherwise noted. See also Supplementary Figure 6 and Supplementary Table 2.

Quantification of the relative proportions of each cell type within tumors showed that HIF2 deletion in αSMA+ CAFs did not affect the total number of fibroblasts within tumors (Supplementary Figure S6). Single-cell analyses were largely concordant with the bulk RNA-seq and IHC data, showing that CAF-HIF2 KO tumors had significantly fewer myeloid cells than CAF-HIF2 WT tumors (14.3% versus 33.1%; n = 3 tumors/group, *P* = 0.045; Figure 2B-D and 4B). Further interrogation of this population revealed higher expression of *Cd11b* (*Itgam*), *Cd68, Adgre1* (F4/80), *Arg1*, and *Mrc1*, indicating a predominance of M2-polarized TAMs (Figure 4C, Supplementary Figure 6E, and Supplementary Table 2). A substantial proportion of these TAMs expressed the immunosuppressive checkpoint ligands *Cd274* (*Pdl1*, ligand for PD-1) and *Cd86* (ligand for CTLA-4; Figure 4D). Single-cell analysis also showed higher expression of *Ctla4, Foxp3*, and *Pdcd1* (*Pd1*) in a subset of T cells (Figures 4E and Supplementary Figure 6E), indicating the presence of Tregs in KPF tumors. Moreover, IHC staining of tumor sections for the Treg marker FoxP3 showed that CAF-HIF2 KO tumors had significantly fewer Tregs than CAF-HIF2 WT tumors (n = 5-6 tumors/group, *P* = 0.014; Figure 4F). These data strongly suggest that deletion of HIF2 in CAFs reduces the PDAC immunosuppressive landscape.

### Inhibition of HIF2 Signaling Enhances PDAC’s Response to Immunotherapy

PDAC is highly resistant to immunotherapy^1^, but recent studies have suggested that targeting non-redundant pathways by combining anti-CTLA4 and anti-PD1 therapies may overcome the inherent TME immunosuppression^30^. Since HIF2 deletion reduced the number of immunosuppressive M2-polarized TAMs and Tregs, we reasoned that PT2399 might improve response to checkpoint immunotherapy. To test this hypothesis, we implanted KPC cells subcutaneously into syngeneic C57BL/6 mice and assigned them to one of four treatments: vehicle plus IgG control, vehicle plus anti-CTLA4 antibody (αCTLA4), PT2399 plus IgG, or PT2399 plus αCTLA4 (Figure 4G). We found that the combination of PT2399 with αCTLA4 significantly slowed tumor growth (n = 10 mice/group, *P =* 0.038), while treatment with either drug alone had no discernible effect (Figure 4H).

We next implanted KPC cells orthotopically into syngeneic C57BL/6 mice to test whether HIF2 inhibition enhanced response to dual checkpoint blockade (DCB) with αCTLA4 and anti-PD1 antibody (αPD1). Mice were assigned to one of four treatments: vehicle plus IgG, vehicle plus DCB, PT2399 plus IgG, or PT2399 plus DCB, with the goal to assess 60-day survival (Figure 4I). The experiment was prematurely terminated due to institutional mandates related to COVID-19, yet the survival rate at 45 days in mice that received combined PT2399 and DCB was 100%, significantly better than the survival rate in the groups treated with IgG control (n = 10 mice/group, *P ≤* 0.0005), and trending toward improved survival compared to mice treated with DCB and vehicle (n = 10 mice/group, *P* = 0.067; Figure 4J). Taken together, these results suggest that HIF2 inhibition might enhance anti-tumor immune responses and improve survival.

## Discussion

Our study addresses a long-standing knowledge gap about the relative roles of HIF signaling in the PDAC microenvironment. Here we show that CAF-specific expression of HIF2, but not HIF1, drives a subset of signals that increases the presence of immunosuppressive cells like TAMs and Tregs in a HIF2-dependent fashion. Furthermore, genetic or pharmacologic inhibition of HIF2 improved survival in spontaneous and syngeneic mouse models.

Our study identifies HIF2 signaling in CAFs as a critical component of hypoxia-related immunosuppression in pancreatic cancer. We demonstrate that HIF2 signaling orchestrates immunosuppression within pancreatic tumors by shifting the cellular composition of the TME, rather than by altering fibrosis, which was unchanged in our model. We observed more TAMs and Tregs in tumors from mice with intact HIF2 function compared to mice with HIF2 deletion in CAFs. These data contrast with findings from a previous study in which depletion of αSMA+ CAFs reduced fibrosis and increased Tregs and cancer progression ^8^. These phenotypic differences are most likely explained by the different approaches of the two studies: ours targeted CAF functionality under hypoxia, while the former study ablated CAFs altogether. We note that we used our dual recombinase system to interrogate only αSMA+ CAFs. Recent studies have shown tumor-supportive roles for other CAF subtypes that do not express αSMA^31^. These other subtypes could be studied in future experiments using different Cre drivers, such as *Fap* or *Fsp1*, to address the dynamic relationships between CAF populations.

While hypoxia, and therefore HIF signaling, affects nearly the entire pancreatic tumor, our data suggest that the detrimental effects of tumor hypoxia are mediated by HIF2 in CAFs, which has not been previously reported. Given that CAFs are the main component and producers of tumor stroma, and that years of accumulating evidence point to hypoxia having a role in PDAC’s aggressiveness and resistance to therapies, our data advances the field’s understanding of how the hypoxic stroma interplays to promote a cold, immune-hostile microenvironment, that is able to overcome immunotherapy. We demonstrate that hypoxic CAFs, reprogrammed in the TME, could stimulate the migration and polarization of macrophages in a HIF2-dependent fashion when cultured *ex vivo*. Importantly, fibroblasts isolated from normal pancreata had no discernible influence on macrophage function. It is not yet known if this novel crosstalk is mediated by a single soluble factor or by a combination of growth factors^27^, metabolites^32^, and/or exosomes^33^. The effects of tumor hypoxia on erratic angiogenesis have been postulated as an underlying mechanism of ineffective tumor infiltration in PDAC^28^. Yet, this mechanism is not consistent with our model, since pharmacological inhibition of HIF2 in CAFs did not alter their expression of *Vegf*, the main driver of tumor angiogenesis^28, 29^. Other groups have postulated roles for cytokines like CSF1, which enhances the recruitment and polarization of macrophages^27^. A potential role of CSF1 is consistent with our sequencing data which demonstrate reduced expression of *Csf1r* in CAF-HIF2 KO tumors. Although studies in carcinogen-induced inflammatory cancer models showed that HIF2 directly regulates macrophage migration and polarization by inducing CSF1R^34^, our data suggest that in pancreatic cancer, HIF2 regulates macrophages indirectly through CAFs. Moreover, our data does not rule out the contribution of HIF2-dependent metabolic^32^ or epigenetic^35^ changes within CAFs.

We demonstrated with single-cell resolution that M2-polarized TAMs were a major source of CD86 and PD-L1 in the TME, whereas their respective receptors were expressed in a subset of T cells. Thus, we infer that these TAMs may be partially responsible for the subsequent reprogramming of effector T cells in the pancreatic TME. Deletion of CAF-HIF2 correlated with reduced TAM density and improved survival from pancreatic cancer in mice, similar to the findings in several clinical reports^26, 27^.

Furthermore, we show that the effects of HIF2 in the TME can be modulated with the clinical HIF2 inhibitor PT2399, which enhanced immune responses in syngeneic models. This drug is already in advanced trials for renal cell carcinoma^22^ and could potentially be repurposed to treat pancreatic cancer as part of a future clinical trial. In summary, this study shows the importance of CAF-specific HIF2 signaling in regulating the PDAC immune landscape and highlights potential novel therapeutic avenues.

## Acknowledgements

We would like to acknowledge Dr. David Kirsch (Duke) and Dr. Dieter Saur for their generous gift of KPF breeders for our colony and Dr. Richard Premont (Case Western) for providing the αSMA^CreERT2^ mice. Experimental design figures were made using BioRender.com. We thank the MD Anderson Research Library Editing Services for their input.

## Author Contributions

Conceptualization, Y.H. and C.M.T.; Data Curation, Y.H., C.J.G.G., D.L., and N.D.N.; Formal Analysis, Y.H., C.J.G.G., N.D.N., J.J.L., and V.B.; Funding Acquisition, C.M.T.; Investigation, Y.H., C.J.G.G., D.L., N.D.N., T.N.F., J.Z., M.Y., A.M.D., J.L.P., A.D., and J.M.M.; Methodology, Y.H. and C.M.T.; Project Administration, Y.H., C.J.G.G., and C.M.T.; Resources, D.S., A.M., and C.M.T.; Software, Y.H., N.D.N., J.J.L., and V.B.; Supervision, C.M.T.; Validation, Y.H., C.J.G.G., and C.M.T.; Visualization, Y.H., C.J.G.G., D.L., N.D.N., J.J.L., N.R.F., M.C.T., and C.M.T.; Writing – Original Draft, C.J.G.G., D.L., N.D.N., and C.M.T.; Writing – Review & Editing, C.J.G.G., D.L., M.C.T., and C.M.T.

