## Supplementary Table 3 & Supplementary Figures 1-6 for "Stromal HIF2 Regulates Immune Suppression in the Pancreatic Cancer Microenvironment"

### Supplemental Data

**Supplementary Table 1.** Normalized counts of bulk RNA-seq in KPF CAF-HIF2 KO tumors, Related to Figure 2.
**Supplementary Table 2.** scRNA-seq Genes Enriched in Cell Populations, Related to Figure 4.
**Supplementary Table 3.** PCR and qRT-PCR Primer List.
**Supplementary Figure 1.** Stromal HIF2 but not HIF1 is critical for PDAC progression.
**Supplementary Figure 2.** Histopathological analyses of PDAC tumors with CAF-specific HIF2 ablation.
**Supplementary Figure 3.** KPF pancreatic cancer cell-specific HIF1 or HIF2 knockout does not affect PDAC progression.
**Supplementary Figure 4.** Stromal HIF2 regulates tumor macrophage recruitment.
**Supplementary Figure 5**. Hypoxic CAFs promote macrophage activation and M2 polarization in a HIF2-dependent paracrine fashion.
**Supplementary Figure 6.** Stromal HIF2 ablation reduces the PDAC immunosuppressive landscape.

| **Supplementary Table 3: PCR and RT-PCR Primer List** | |
|---|---|
| Arg1 F | ACAAGACAGGGCTCCTTTCAG |
| Arg1 R | CTTGGGAGGAGAAGGCGTTT |
| C3ar1 F | TGCTCAGCAACTCGTCCAAT |
| C3ar1 F | ATGGAGGCAATGTCTTGGGG |
| Cd74 F | CTCCTTGGGCCTGTGAAGAA |
| Cd74 R | GTTACCGTTCTCGTCGCACT |
| Hif2a F | CAGGCAGTATGCCTGGCTAATTCCAGTT |
| Hif2a R-flox | CTTCTTCCATCATCTGGGATCTGGGACT |
| Hif2a R-KO | GCTAACACTGTACTGTCTGAAAGAGTAGC |
| Itgam F | GGCAGCCAGATTGGCTCTTA |
| Itgam R | GCTTCACACTGCCACCGT |
| Mmp9 F | CAGCCGACTTTTGTGGTCTTC |
| Mmp9 R | GTACAAGTATGCCTCTGCCA |
| Tgfb-1 F | ACCGCAACAACGCCATCTAT |
| Tgfb-1 R | TGCCGTACAACTCCAGTGAC |
| Vegfa F | TTCGTCCAACTTCTGGGCTC |
| Vegfa R | CTGGGACCACTTGGCATGG |

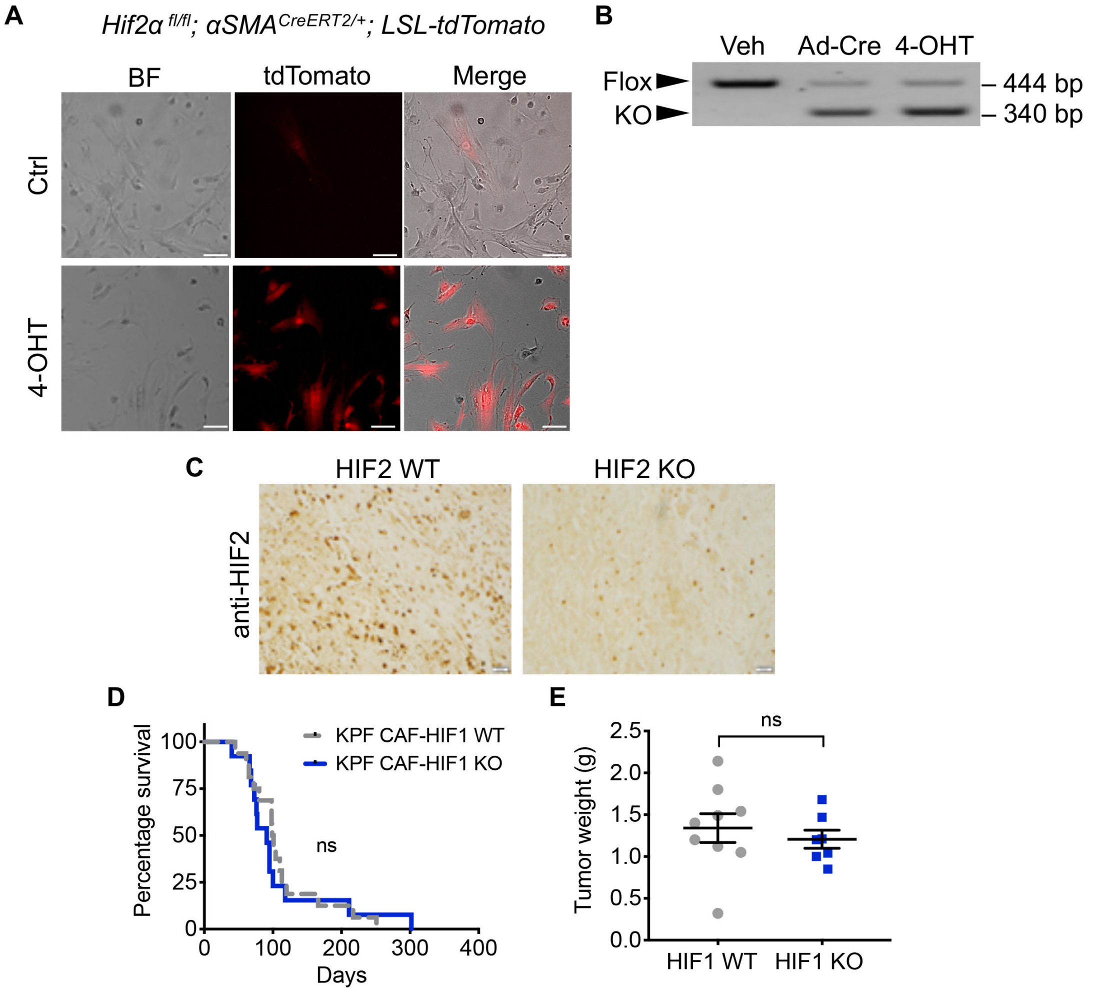

**Supplementary Figure 1. Stromal HIF2 but not HIF1 is critical for PDAC progression, Related to Figure 1.** *(A and B)* Fibroblasts isolated from the pancreata of *Hif2α$^{fl/fl}$; αSMA$^{CreERT2/+}$; LSL-tdTomato* mice were treated with vehicle DMSO (veh), adenovirus-Cre (Ad-Cre) as a positive control, or 4-hydroxytamoxifen (4-OHT) *ex vivo*. *(A)* Fluorescent microscopy confirmed activation of the Cre-ERT2 fusion protein in activated αSMA+ fibroblasts. *(B)* PCR genotyping confirmed deletion of HIF2. Scale bars, 100 μm; BF, bright-field images. *(C)* Representative IHC images of KPF CAF-HIF2 WT and KO tumors stained for HIF2 (n = 5/group). Scale bars, 20 μm. *(D)* Kaplan-Meier curves showing percentage survival for KPF CAF-HIF1 WT (n = 16) and KO mice (n = 13). ns (not significant), by log-rank test. *(E)* Corresponding tumor weights for KPF CAF-HIF1 WT (n = 9) and KO mice (n = 7). Mean ± SEM; ns, by Student's *t* test.

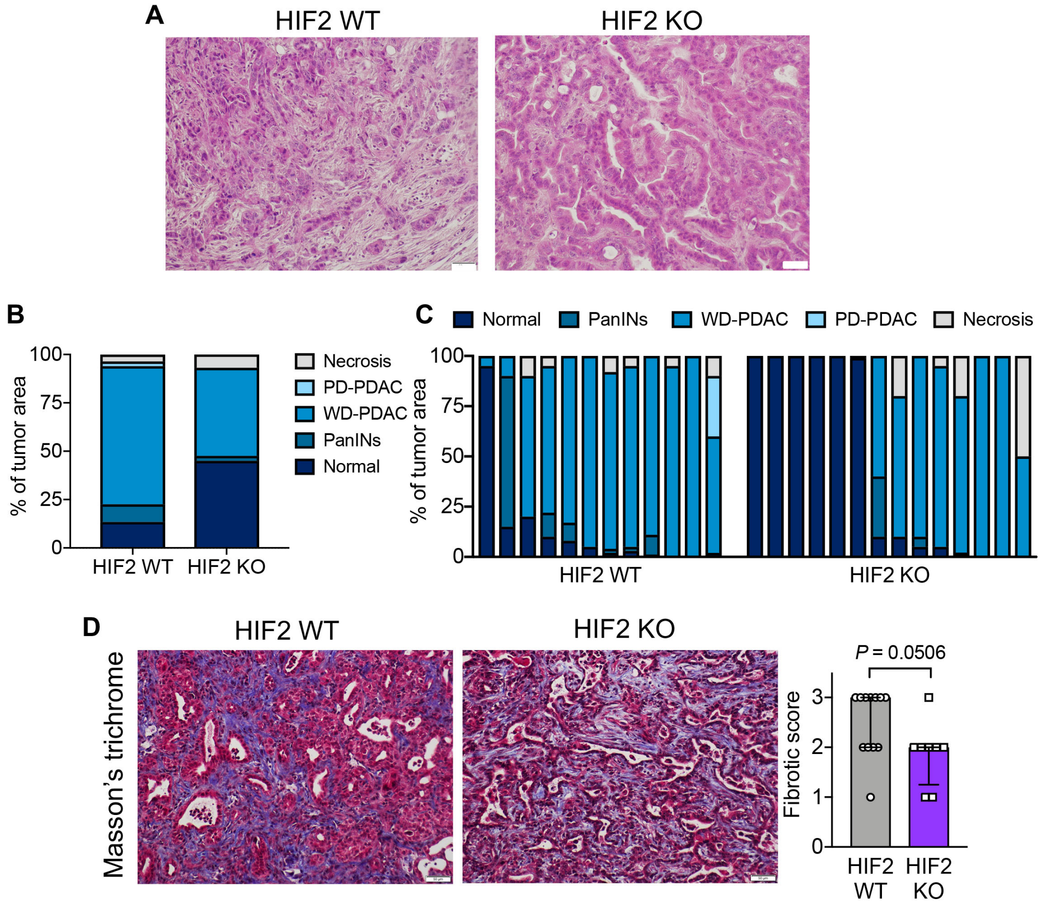

**Supplementary Figure 2. Histopathological analyses of PDAC tumors with CAF-specific HIF2 ablation, Related to Figure 1.** *(A)* Representative images of H&E-stained KPF CAF-HIF2 WT (n = 12) and KO (n = 14) tumors. Scale bars, 50 μm. *(B and C)* Quantification of the differentiation state of the tumors in *(A)*. The fractions of each pancreas that were necrotic tissue, poorly differentiated PDAC (PD-PDAC), well-differentiated PDAC (WD-PDAC), pancreatic intraepithelial neoplasia (PanIN), or normal tissue were scored in a blinded manner, and values were averaged and compared between groups *(B)* or shown individually *(C)*. *(D) Left:* Representative images of Masson's trichrome staining in KPF CAF-HIF2 WT (n = 12) and KO (n = 8) tumors; scale bars, 50 μm. *Right:* Fibrotic score; median ± interquartile range; *P*, by Mann–Whitney *U* test.

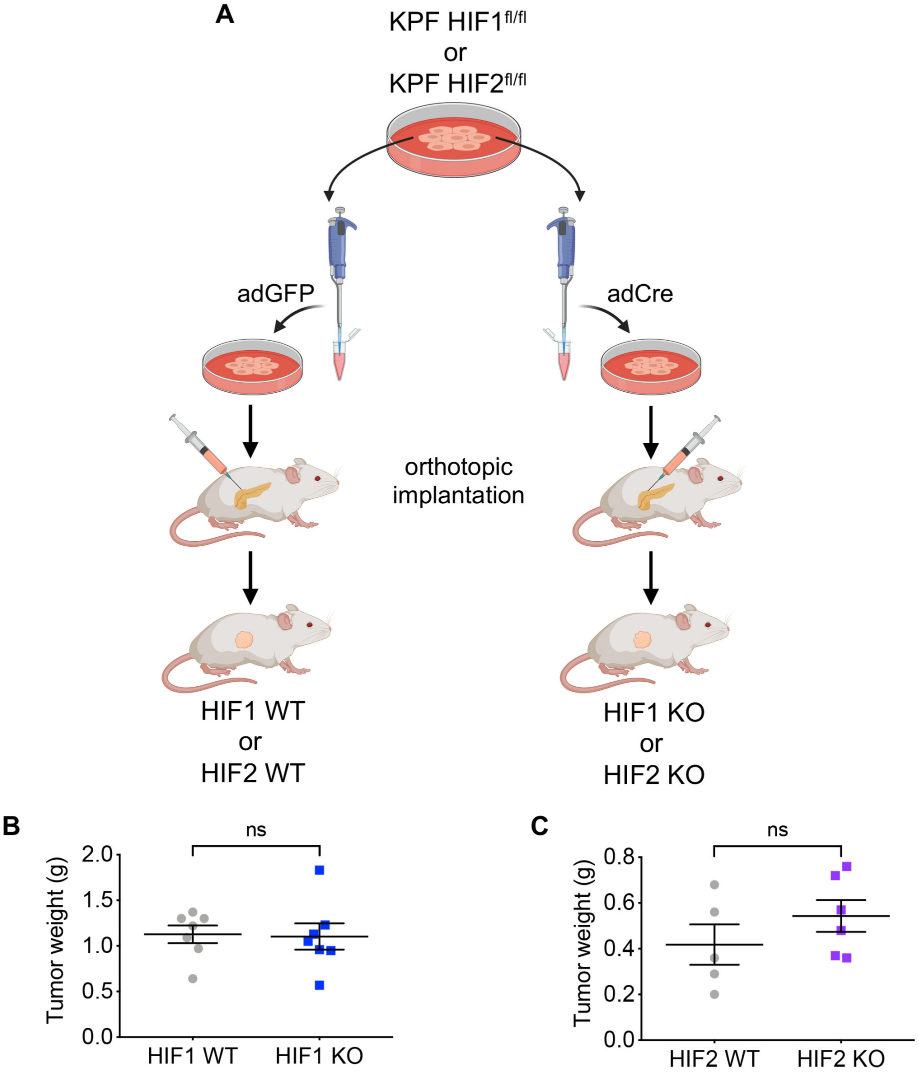

**Supplementary Figure 3. KPF pancreatic cancer cell-specific HIF1 or HIF2 knockout does not affect PDAC progression, Related to Figure 1.**
*(A)* Cancer cells from KPF *Hif1a*$^{fl/fl}$ or *Hif2a*$^{fl/fl}$ tumors were isolated and infected *ex vivo* with GFP adenovirus (adGFP) or Cre adenovirus (adCre) and subsequently injected into the pancreata of immunocompromised mice. *(B and C)* Comparison of the orthotopic tumors showed no significant difference in tumor size between tumors with *(B) Hif1a* or *(C) Hif2a* deletion and the respective WT controls. Mean ± SEM; ns, by Student's *t* test.

A

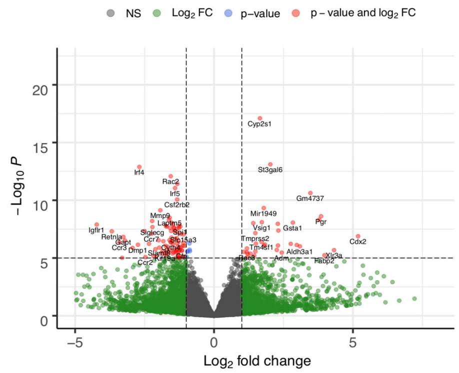

B

Enriched in KPF CAF-HIF2 WT tumors (compared to KPF CAF-HIF2 KO tumors)

| Rank | Gene set | MSigDB Collection | NES | Nom p-value | FDR q-value |
|---|---|---|---|---|---|
| 1 | REACTOME_DAP12_SIGNALING | C2 | 2.15 | 0.000 | 0.014 |
| 2 | REACTOME_INTEGRIN_SIGNALING | C2 | 2.06 | 0.000 | 0.023 |
| 3 | KEGG_FC_GAMMA_R_MEDIATED_PHAGOCYTOSIS | C2 | 1.98 | 0.000 | 0.020 |
| 4 | GO_REGULATION_OF_LYMPHOCYTE_MEDIATED_IMMUNITY | C5 | 1.88 | 0.000 | 0.091 |
| 5 | GO_MYELOID_CELL_APOPTOTIC_PROCESS | C5 | 1.85 | 0.000 | 0.088 |
| 6 | HALLMARK_KRAS_SIGNALING_UP | H | 1.83 | 0.000 | 0.040 |
| 7 | GO_REGULATION_OF_MYELOID_LEUKOCYTE_MEDIATED_IMMUNITY | C5 | 1.81 | 0.034 | 0.077 |
| 8 | HALLMARK_INFLAMMATORY_RESPONSE | H | 1.80 | 0.000 | 0.043 |
| 9 | GO_RESPIRATORY_BURST | C5 | 1.79 | 0.000 | 0.081 |
| 10 | GO_MACROPHAGE_ACTIVATION | C5 | 1.78 | 0.000 | 0.084 |
| 11 | GO_POSITIVE_REGULATION_OF_INTERLEUKIN_12_PRODUCTION | C5 | 1.76 | 0.000 | 0.084 |
| 12 | GO_MACROPHAGE_DIFFERENTIATION | C5 | 1.76 | 0.000 | 0.085 |
| 13 | GO_POSITIVE_REGULATION_OF_TUMOR_NECROSIS_FACTOR_SUPERFAMILY_CYTOKINE_ PRODUCTION | C5 | 1.75 | 0.000 | 0.085 |
| 14 | GO_INTERLEUKIN_10_PRODUCTION | C5 | 1.74 | 0.000 | 0.090 |
| 15 | GO_DENDRITIC_CELL_MIGRATION | C5 | 1.71 | 0.000 | 0.098 |
| 16 | GO_POSITIVE_REGULATION_OF_INTERLEUKIN_6_PRODUCTION | C5 | 1.69 | 0.032 | 0.104 |
| 17 | GO_POSITIVE_REGULATION_OF_INTERLEUKIN_4_PRODUCTION | C5 | 1.65 | 0.000 | 0.127 |
| 18 | GO_INTERLEUKIN_1_BETA_PRODUCTION | C5 | 1.62 | 0.000 | 0.145 |
| 19 | GO_NEGATIVE_REGULATION_OF_CELLULAR_RESPONSE_TO_TRANSFORMING_GROWTH_ FACTOR_BETA_STIMULUS | C5 | 1.61 | 0.000 | 0.085 |
| 20 | GO_REGULATION_OF_MACROPHAGE_MIGRATION | C5 | 1.60 | 0.020 | 0.161 |

C

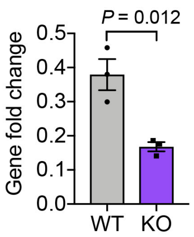

**Supplementary Figure 4. Stromal HIF2 regulates tumor macrophage recruitment, Related to Figure 2 and Supplementary Table 1.** *(A)* Volcano plot illustrating differential gene expression analysis using bulk RNA-seq data from KPF CAF-HIF2 tumors (n = 4/group). *(B)* Gene sets enriched in KPF CAF-HIF2 WT tumors compared to KO tumors ranked by normalized enrichment score (NES). Gene set enrichment analysis was performed using bulk RNA-seq data from *(A)*. Nom., nominal; FDR, false discovery rate. *(C)* qRT-PCR confirmed the downregulation of *C3ar1* in CAF-HIF2 KO tumors. Error bars represent mean ± SEM, and each dot represents 1 tumor sample. *P*, by Student's *t* test.

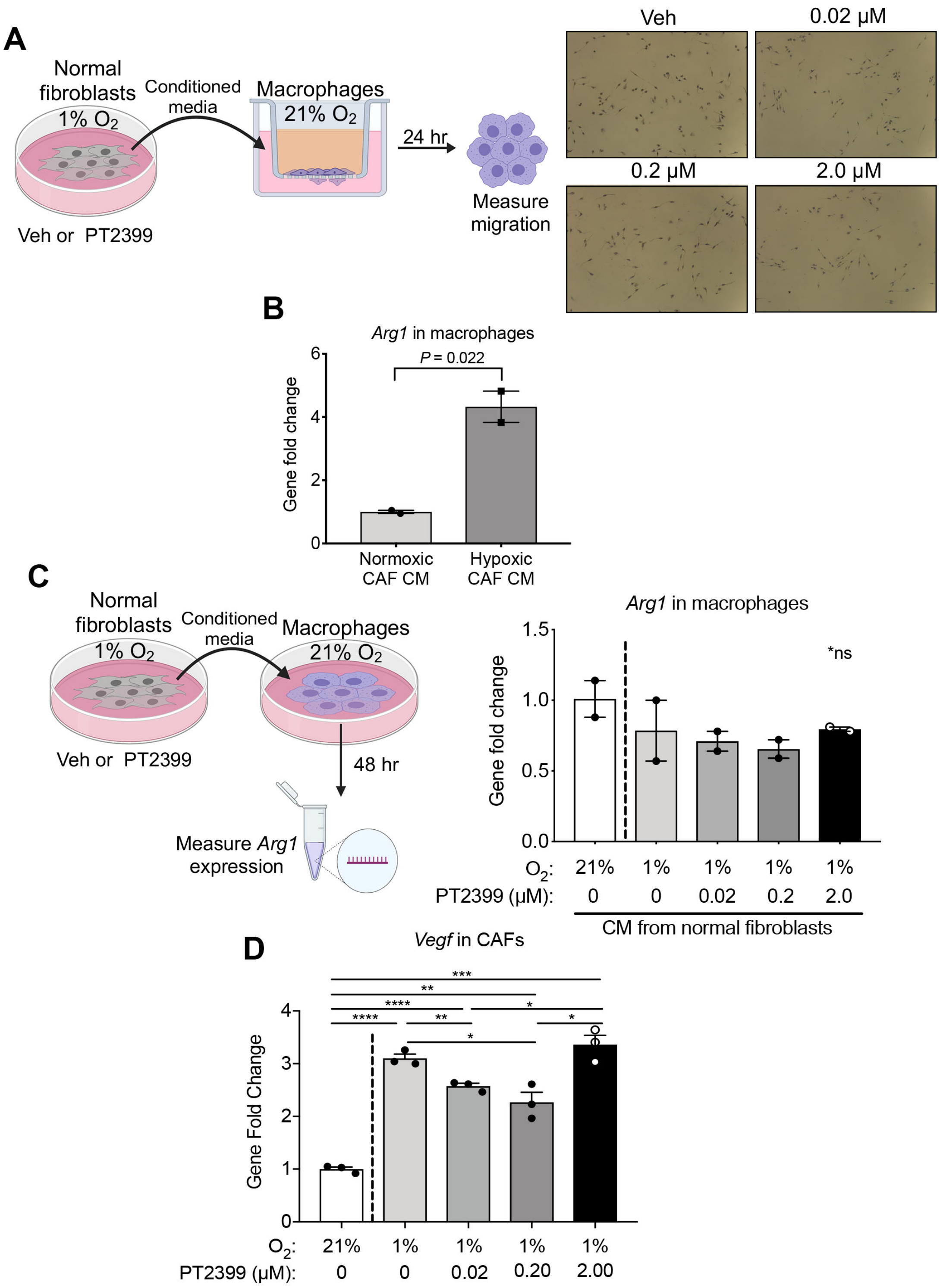

**Supplementary Figure 5. Hypoxic CAFs promote macrophage activation and M2 polarization in a HIF2-dependent paracrine fashion, Related to Figure 3.**
*(A)* Macrophages were cultured in transwell plates and incubated with conditioned media (CM) collected from hypoxic normal fibroblasts treated with vehicle (veh) or PT2399. Representative bright-field images of the transwell assay are shown. *(B)* Macrophages were incubated with CM collected from CAFs grown in normoxic or hypoxic conditions and

*Arg1* expression was measured by qRT-PCR. *(C)* Macrophages were incubated with CM collected from hypoxic normal fibroblasts treated with vehicle or PT2399 and *Arg1* expression was measured by qRT-PCR. *(D)* CAFs were grown in normoxic or hypoxic conditions and treated with PT2399 at the indicated doses, then *Vegfa* expression was measured by qRT-PCR. All error bars represent mean ± SEM; *P*, by Student's *t* test; **P* ≤ .05, ***P* ≤ .01, ****P* ≤ .001, *****P* ≤ .0001; ns, not significant.

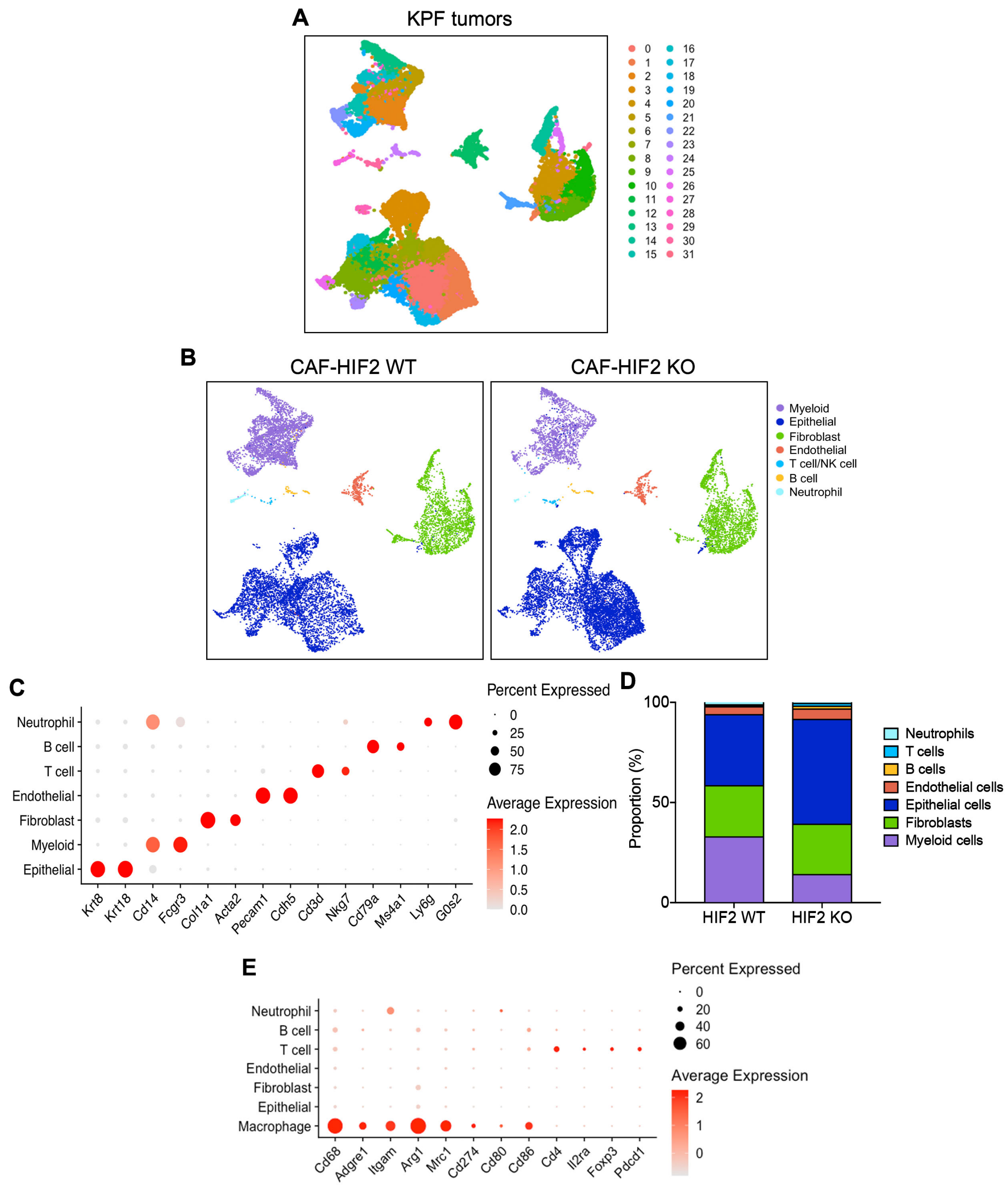

**Supplementary Figure 6. Stromal HIF2 ablation reduces the PDAC immunosuppressive landscape, Related to Figure 4 and Supplementary Table 2.** *(A)* UMAP of scRNA-seq analysis of 22,635 cells sorted from 6 mice (3/group). Graph-based clustering identified 32 clusters, indicated by color. *(B)* UMAP of scRNA-seq analysis of 22,635 cells isolated from KPF CAF-HIF2 WT tumors (left; 10,703 cells) and KO tumors (right; 11,932 cells). *(C)* Bubble plots showing the relative average expression of selected cell-type-specific markers across all major cell populations identified in the scRNA-seq analysis. The size of the dots indicates the percentage of the cell population that expressed the marker, and the intensity of color indicates the average expression level. *(D)* Proportions of cell types in CAF-HIF2 WT and KO tumors, quantified as an average per group. *(E)* Bubble plots showing the relative average expression of macrophage and Treg markers across all major cell populations identified in the scRNA-seq analysis. The size of the dots indicates the percentage of the cell population that expressed the marker, and the intensity of color indicates the average expression level.
